# *Toxoplasma gondii* AP2XII-2 contributes to proper progression through S-phase of the cell cycle

**DOI:** 10.1101/2020.06.09.143586

**Authors:** Sandeep Srivastava, Michael W. White, William J. Sullivan

**Affiliations:** Department of Pharmacology & Toxicology, Indiana University School of Medicine, Indianapolis, 46202, Indiana; Department of Microbiology & Immunology, Indiana University School of Medicine, Indianapolis, 46202, Indiana; Department of Global Health, University of South Florida, Tampa, Florida

## Abstract

*Toxoplasma gondii* is a protozoan parasite that causes lifelong chronic infection that can reactivate in immunocompromised individuals. Upon infection, the replicative stage (tachyzoite) converts into a latent tissue cyst stage (bradyzoite). Like other apicomplexans, *T. gondii* possesses an extensive linage of proteins called ApiAP2s that contain plant-like DNA-binding domains. The function of most ApiAP2s is unknown. We previously found that AP2IX-4 is a cell cycle-regulated ApiAP2 expressed only in dividing parasites as a putative transcriptional repressor. In this study, we purified proteins interacting with AP2IX-4, finding it to be a component of the recently characterized microrchidia (MORC) transcriptional repressor complex. We further analyzed AP2XII-2, another cell cycle-regulated factor that associates with AP2IX-4. We monitored parallel expression of AP2IX-4 and AP2XII-2 proteins in tachyzoites, detecting peak expression during S/M phase. Unlike AP2IX-4, which is dispensable in tachyzoites, loss of AP2XII-2 resulted in a slowed tachyzoite growth due to a delay in S-phase progression. We also found that AP2XII-2 depletion increased the frequency of bradyzoite differentiation in vitro. These results suggest that multiple AP2 factors collaborate to ensure proper cell cycle progression and tissue cyst formation in *T. gondii*.

**Importance:** *Toxoplasma gondii* is a single-celled parasite that causes opportunistic infection due to its ability to convert into a latent cyst stage. This work describes a new transcriptional factor called AP2XII-2 that plays a role in properly maintaining the growth rate of replicating parasites, which contributes to signals required for development into its dormant stage. Without AP2XII-2, *Toxoplasma* parasites experience a delay in their cell cycle that increases the frequency of latent cyst formation. In addition, we found that AP2XII-2 operates in a multi-subunit complex with other AP2 factors and chromatin remodeling machinery that represses gene expression. These findings add to our understanding of how *Toxoplasma* parasites balance replication and dormancy, revealing novel points of potential therapeutic intervention to disrupt this clinically relevant process.

## Introduction

*Toxoplasma gondii*, an obligate intracellular apicomplexan parasite of medical and veterinary interest, can infect almost all warm-blooded animals and is present in one-third of the human population. The complex life cycle of *T. gondii* consists of multiple warm-blooded hosts and different developmental forms, involving asexual and sexual stages. The sexual cycle takes place exclusively in the gut of felines, the definitive host for the life cycle, which consequently excrete infectious oocysts into the environment (1). The asexual phase of the life cycle is comprised of replicating tachyzoites and quiescent bradyzoites. Upon infection, the parasite increases its biomass and disseminates throughout the body as tachyzoites, which subsequently convert into latent bradyzoites that persist in the brain, heart, and skeletal muscle inside intracellular tissue cysts. The presence of bradyzoite cysts in the meat and organs of infected animals is another major route of *T. gondii* transmission (2).

Despite the high prevalence in humans, acute toxoplasmosis is rarely seen, most commonly observed as a reactivated infection in patients with HIV/AIDS or some other immunocompromised condition (3,4). Congenital toxoplasmosis can also occur if tachyzoites traverse the placenta, which can lead to miscarriage or birth defects in humans, as well as abortion in livestock (5-7). Bradyzoites tissue cysts do not appear to be cleared effectively by the immune system of the host, nor are they targeted by the currently approved therapeutics (8-10). The formation of latent bradyzoite cysts is central to *T. gondii* pathogenesis and transmission, but the molecular mechanisms involved in stage conversion are incompletely understood (11,12).

The hunt for *T. gondii* transcription factors that coordinate the reprogramming of the genome required for stage conversion has been challenging due to a striking lack of conventional master regulators in the parasite genome (13). In 2005, a new family of Apicomplexa proteins that possess a DNA-binding domain related to the Apetala-2 transcription factors of plants was identified (14). These proteins are called ApiAP2 factors and the *T. gondii* genome encodes 67 of these proteins (15-17), while the fellow apicomplexan parasite *Plasmodium falciparum* has 27 (14). In each parasite, multiple ApiAP2s have been linked to playing a role in regulating gene expression during specific developmental stages. For example, *Plasmodium* AP2-G regulates gametocyte development (18) and AP2-Sp2 regulates gene expression during the sporozoite stage (19). A number of ApiAP2s in *T. gondii* have been linked to tachyzoite stage conversion into bradyzoites (20). AP2IX-9 acts as a transcriptional repressor of bradyzoite genes and restricts commitment to develop into in vitro bradyzoite tissue cysts (20). Like AP2IX-9, AP2IV-3 is upregulated during pH-induced in vitro bradyzoite differentiation, but acts as a transcriptional activator that likely competes to control bradyzoite gene expression with AP2IX-9 (21). Knocking out AP2IV-4 resulted in the expression of a subset of bradyzoite-specific proteins in replicating tachyzoites, which prevented tissue cyst formation in mice (22).

We previously determined that AP2IX-4 is a cell-cycle regulated factor expressed exclusively during division of tachyzoites and bradyzoites (23). Genetic knockout of AP2IX-4 had no discernible effect on tachyzoites, but reduced the frequency of tissue cyst formation in vitro and in vivo in mice (23). Transcriptional profiling showed that PruΔ*ap2IX-4* parasites display upregulation of a subset of bradyzoite genes over and above that seen in parental parasites exposed to alkaline stress, suggesting that it functions as a repressor of these genes. Despite the upregulation of several key bradyzoite genes, PruΔ*ap2IX-4* parasites showed a modest decrease in bradyzoite conversion in culture and in BALB/c mice.

To better understand the role of AP2IX-4, we purified the AP2IX-4 complex in this study, finding that it associates with the microrchidia (MORC) transcriptional repressor complex (24). We further interrogated the function of an AP2IX-4-interacting protein, AP2XII-2, which largely shares its cell cycle expression profile. In contrast to the dispensable AP2IX-4, tachyzoites lacking AP2XII-2 exhibit a fitness defect in vitro. Further experiments show that AP2XII-2 is important for proper progression through the S-phase of the tachyzoite cell cycle and that its loss increases in vitro bradyzoite differentiation. These findings shed new light on the complex interplay between multiple AP2 factors regulating cell cycle and bradyzoite development in *T. gondii*.

## Results

### Identification of proteins interacting with AP2IX-4

We previously identified a cell-cycle regulated ApiAP2 transcription factor called AP2IX-4 (TGME49_288950) that is expressed exclusively in dividing parasites (23). Genetic depletion of AP2IX-4 resulted in dysregulation of stage-specific gene regulation, which lowered the frequency of bradyzoite tissue cyst formation in vitro and in vivo. To better understand the role of AP2IX-4, we engineered RHΔ*ku80* tachyzoites to express endogenous AP2IX-4 protein with a C-terminal 3xHA tag for use in immunoprecipitation experiments (AP2IX-4^HA^) (Fig. 1A). Western analysis of fractionated nuclear and cytosolic extracts revealed that AP2IX-4 is exclusively expressed in the parasite nucleus (Fig. 1B). AP2IX-4^HA^ protein of the expected size (∼104 kDa) is immunoprecipitated using α-HA and nuclear fractions made from purified intracellular parasites (Fig. 1C).

**Figure 1.**
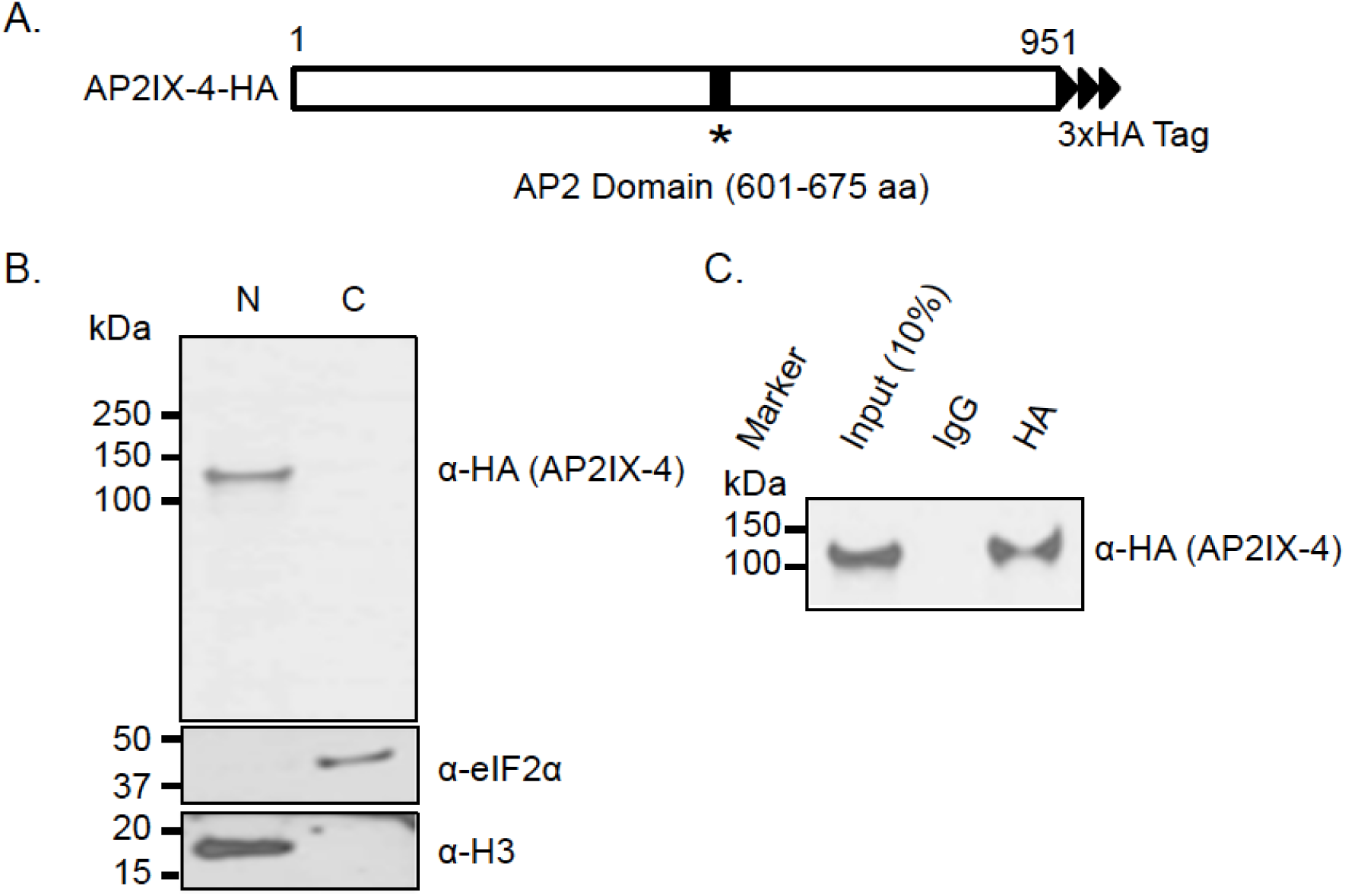
Endogenously tagged AP2IX-4^HA^ localizes to parasite nucleus. **(A)** Diagram of *T. gondii* AP2IX-4^HA^ protein. **(B)** Nuclear (N) and cytosolic (C) fractions of RHΔ*ku80* tachyzoites were generated for western blot analysis using rat α-HA antibodies. Blot was re-probed with α-TgIF2α as a cytosolic marker and α-histone H3 as a nuclear marker. **(C)** Western blot of AP2IX-4 immunoprecipitated from purified, intracellular RHΔ*ku80* tachyzoites via Dynabeads coupled with mouse α-HA antibodies. Western blot was probed using rat α-HA antibody. Tagged AP2IX-4^HA^ migrates at the expected size of ∼104 kDa.

Mass spectrometry and SAINT analysis of proteins co-immunoprecipitating with AP2IX-4^HA^ included two additional ApiAP2 factors: AP2XII-2 (TGGT1_217700) and AP2VIIa-3 (TGGT1_205650) (Fig. 2A; see Supplemental Table 1 for complete dataset). We also detected proteins interacting with AP2IX-4 that are associated with transcriptional repressive activity in tachyzoites: HDAC3 (TGGT1_227290) and CRC230 (TGGT1_305340), and two hypothetical proteins (TGGT1_214140) and (TGGT1_275680) (Fig. 2A). Together, the proteins complexed with AP2IX-4 overlap the recently elucidated microrchidia (MORC) repressor complex that has been implicated in silencing genes associated with the *T. gondii* sexual stages (24). Thus, our findings bolster the observation that multiple AP2 factors recruit MORC to regulate gene expression, and provides an explanation as to why the loss of AP2IX-4 resulted in the dysregulation of genes (23).

**Figure 2.**
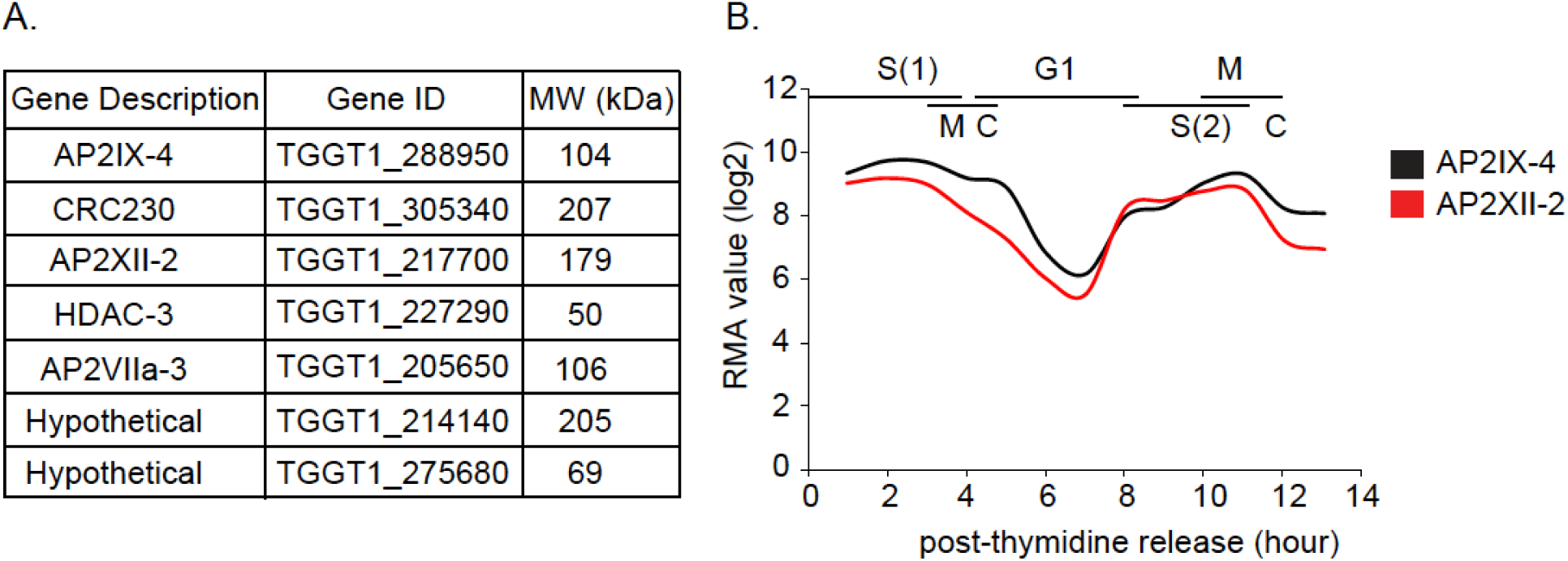
AP2IX-4 core complex in *Toxoplasma* tachyzoites. **(A)** Mass spectrometry results of endogenously tagged AP2IX-4^HA^ immunoprecipitate from tachyzoites, performed on intracellular parasites harvested from infected HFFs. Listed are proteins interacting with AP2IX-4 with probability of interaction as measured by pSAINT of 0.9 or above (see Supplemental Table 1 for complete dataset). **(B)** Expression of transcripts for AP2IX-4 and AP2XII-2 (data source: ToxoDB.org).

Figure 2B shows that mRNA expression profiles of AP2IX-4 and AP2XII-2 have nearly parallel cell cycle-dependent expression patterns (based on data in ref. 22). The other interacting proteins exhibited no cell cycle regulation. Both AP2 mRNAs reach their maximum expression during the S/M phase of the cell cycle before dropping in G1 phase. In contrast, all of the other proteins in the AP2IX-4 complex show constitutive expression throughout the cell cycle. These data suggest that AP2IX-4 and AP2XII-2 operate with MORC only during S/M.

### Localization and interaction between AP2IX-4 and AP2XII-2

The single exon of the AP2XII-2 gene encodes a 195 kDa protein containing 1,431 amino acids and a single AP2 domain proximal to the C-terminus (Fig. 3A). To validate the protein-protein interaction and study AP2XII-2 further, we endogenously tagged the C-terminus of AP2XII-2 with 3xMYC in the RHΔ*ku80* parasites expressing AP2IX-4^HA^. Immunoblots of these dual-tagged parasites confirmed the presence of both in the nuclear fraction (Fig. 3B). We performed reciprocal co-immunoprecipitations of AP2IX-4^HA^ and AP2XII-2^MYC^ followed by western blotting, which validated their interaction in intracellular tachyzoites (Fig. 3C,D).

**Figure 3.**
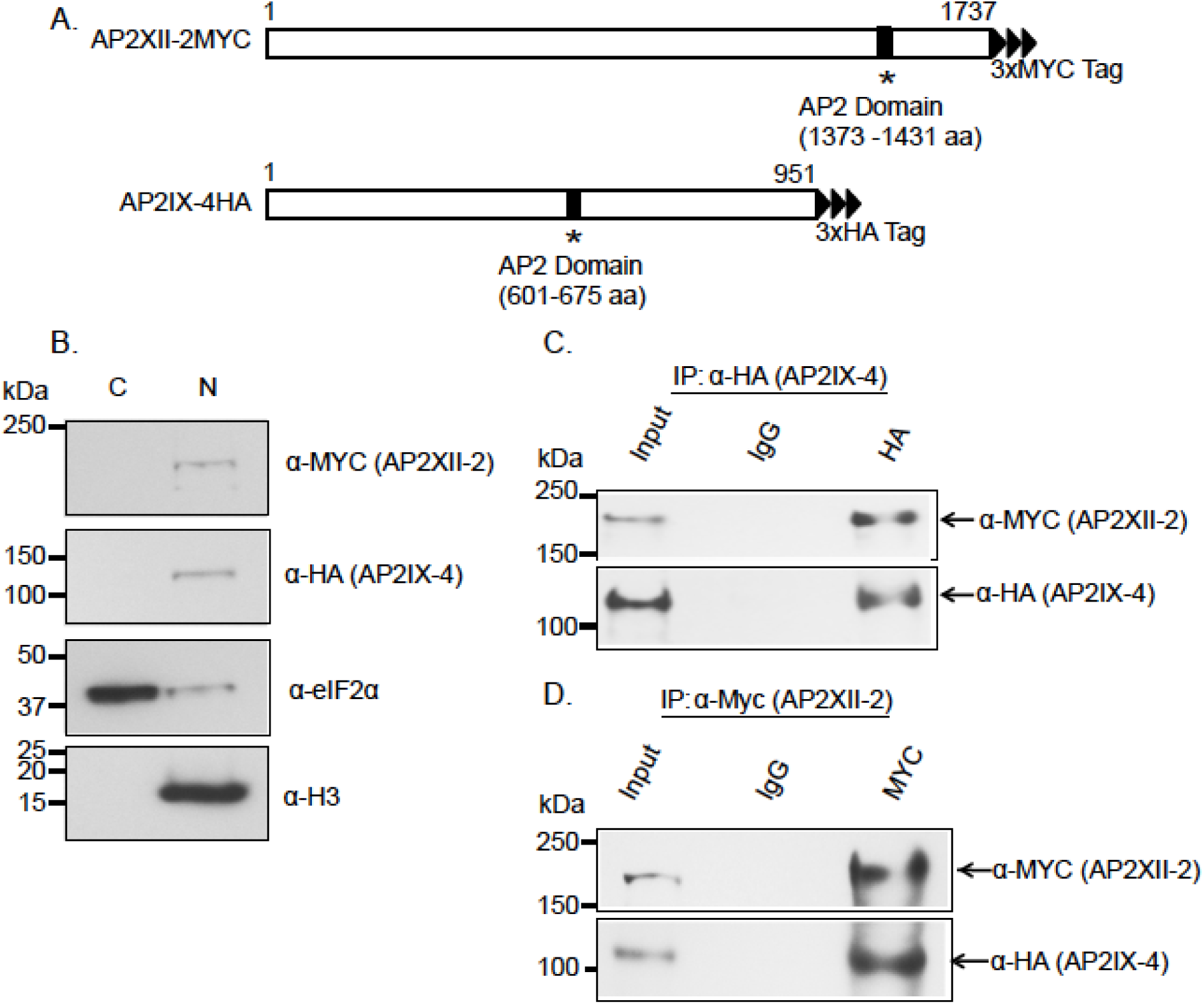
Interaction of AP2IX-4 and AP2XII-2. **(A)** Schematic representation of candidate interacting protein AP2XII-2 endogenously tagged with a c-MYC epitope. **(B)** Western blot of fractionated parasites shows that AP2XII-2^MYC^ and AP2IX-4^HA^ both localize to parasite nucleus. Blot was re-probed with α-TgIF2α and α-histone H3 as cytosolic and nuclear markers, respectively. **(C)** Co-immunoprecipitation confirms the interaction between AP2XII-2^MYC^ and AP2IX-4^HA^ in tachyzoites. IP was performed with α-HA to pull down AP2IX-4^HA^; top panel probed with α-MYC detects AP2XII-2^MYC^ in the IP. Bottom panel shows the membrane re-probed with α-HA. **(D)** Reciprocal co-IP further confirms the interaction: in this case, the IP was performed with α-MYC to pull down AP2XII-2^MYC^; top panel probed with α-MYC detects AP2XII-2^MYC^. Bottom panel shows re-probing with α-HA, which detects AP2IX-4^HA^ in the IP.

We next monitored the pattern of expression for AP2XII-2^MYC^ by immunofluorescence assay (IFA) for comparison to what we observed previously for AP2IX-4 (23). To delineate the cell cycle expression profile of AP2XII-2, we co-stained with established cell cycle markers including Inner membrane complex-3 (IMC3) and TgCentrin-1 (Fig. 4A) (25,26). Results show that AP2XII-2^MYC^ protein is expressed in the second half of the cell cycle prior to the start of budding and continues until buds are mature, in agreement with mRNA profiling (Fig. 2B). This expression pattern is consistent with a cell cycle timing that begins in S phase and continues into mitosis, with the factor downregulated after nuclear division (very early G1 period). Expression of AP2XII-2 diminishes before the onset of C-phase (cytokinesis); this is in contrast to AP2IX-4, which can be detected through to the end of cytokinesis (23). By co-staining with TgCentrin-1, we identified distinct parasite vacuoles in either G1 or S-phase of the cell cycle in the same field; note that AP2XII-2 is only detected in parasites during S-phase and not in G1 (Fig. 4B). By staining with anti-MYC, anti-HA, and anti-IMC3, we were able to show that both AP2 proteins are co-expressed during S-phase of the cell cycle (Fig. 4C). Altogether, these data support an intimate interaction between AP2IX-4 and AP2XII-2 during parasite division.

**Figure 4.**
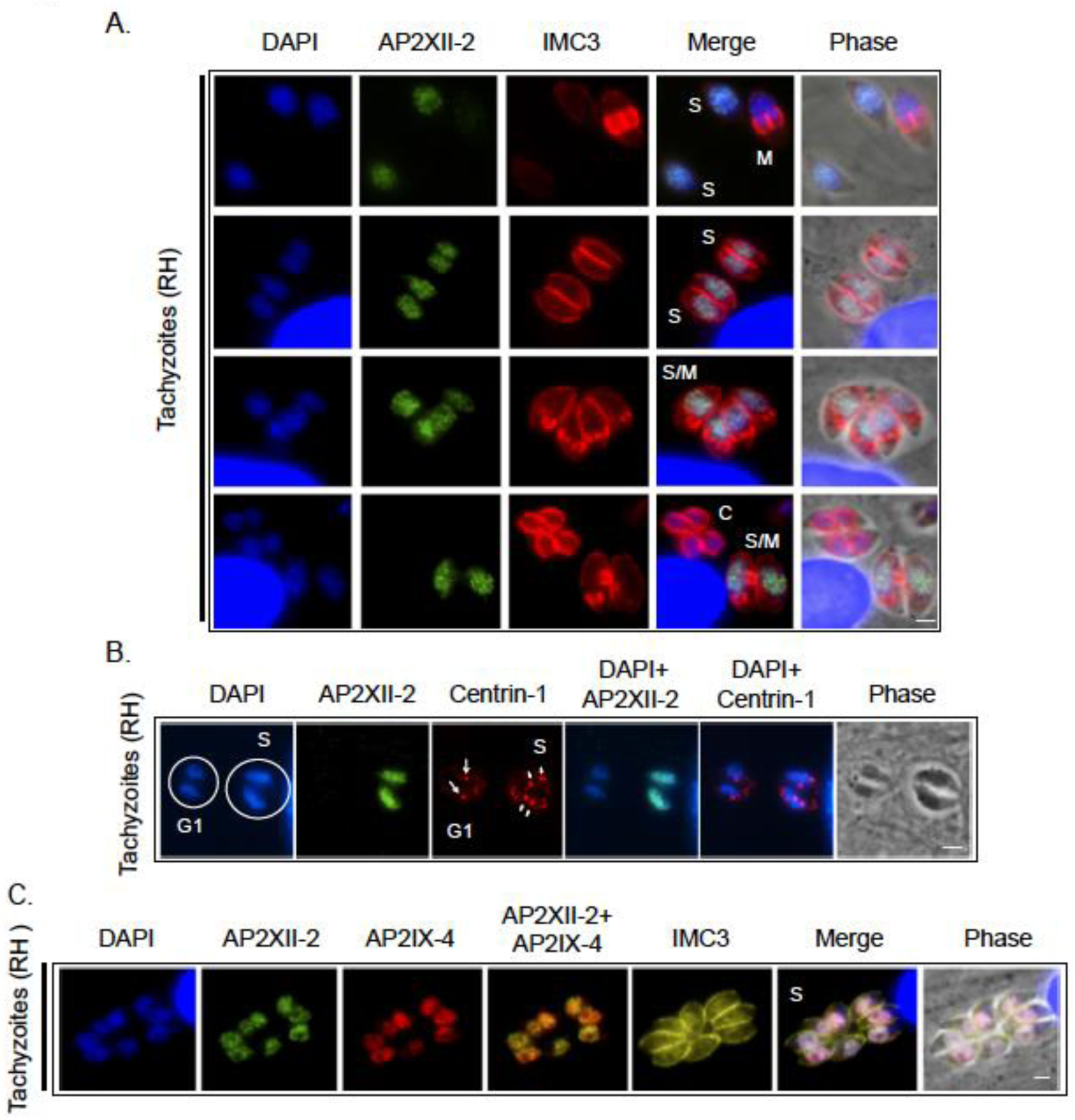
Immunofluorescence assay of AP2IX-4 and AP2XII-2 during cell cycle. **(A)** RHΔ*ku80* parasites expressing endogenously tagged AP2XII-2^MYC^ were probed with antibodies to c-MYC (green) and IMC3 (red), the latter of which helps detect budding daughter parasites. AP2XII-2^MYC^ is detectable in S and S/M phases, but disappears before the onset cytokinesis (C). DAPI (blue) was used as a nuclear stain. The four rows show parasites in different stages of the cell cycle. Scale bar = 2 µM. **(B)** To more closely examine the expression of AP2XII-2^MYC^ during cell cycle, an IFA was performed using antibodies to c-MYC (green) and Centrin-1 (red). Parasites in G1 contain single centrosomes, whereas those in S-phase are duplicated. Arrows indicate centrosomes. Scale bar = 2 µM. **(C)** Triple-labelled IFA to determine cell cycle expression patterns for each AP2 factor. AP2XII-2^MYC^ (green) and AP2IX-4^HA^ (red) co-localize in S-phase. IMC3 (yellow) and larger nuclei (DAPI, blue) identify parasites in S-phase. Scale bar = 2 µM.

### Conditional knockdown of AP2XII-2 slows parasite replication

To further investigate the role of AP2XII-2 in *T. gondii*, we attempted to genetically ablate the endogenous AP2XII-2 gene. Both conventional allelic replacement using homologous recombination as well as CRISPR/Cas9 approaches yielded no viable parasites, suggesting that AP2XII-2 is essential in tachyzoites. Our inability to knockout AP2XII-2 is consistent with a genome-wide CRISPR survey that assigned this AP2 a fitness score of -1.16 (22).

We therefore pursued an IAA-inducible degradation (AID) strategy that would allow a conditional knockdown of AP2XII-2 protein through the addition of IAA (27). We endogenously tagged AP2XII-2 with a C-terminal AID-3xHA tag in RH TIR1-3xFlag *Δku80* parasites (27) (Fig. 5A-B). A clone was isolated in which AP2XII-2^AID-HA^ protein was undetectable within 15 minutes after inclusion of IAA in the culture medium. (Fig. 5C). To determine the role of AP2XII-2 on parasite viability, we performed a standard plaque assay in the presence of IAA. Compared to parental parasites, the loss of AP2XII-2 resulted in smaller plaque sizes, although there was no change in plaque number (Fig. 5D, E). These results suggest a defect in parasite replication rather than invasion. To confirm this idea, we cultured parasites in 500 µM IAA for 24 hours, then syringe-lysed and filter-purified them for an attachment and invasion assay (28). We found that AP2XII-2 depletion had no effect on invasion or attachment of parasites (Fig. 5F). We also conducted a parasite counting assay in the presence of IAA or vehicle, finding that replication rates were significantly lower without AP2XII-2 (Fig. 5G). AP2XII-2^AID-HA^ parasites treated with IAA for 24 hours contained more vacuoles with only 8 parasites and fewer vacuoles with 16 parasites.

**Figure 5.**
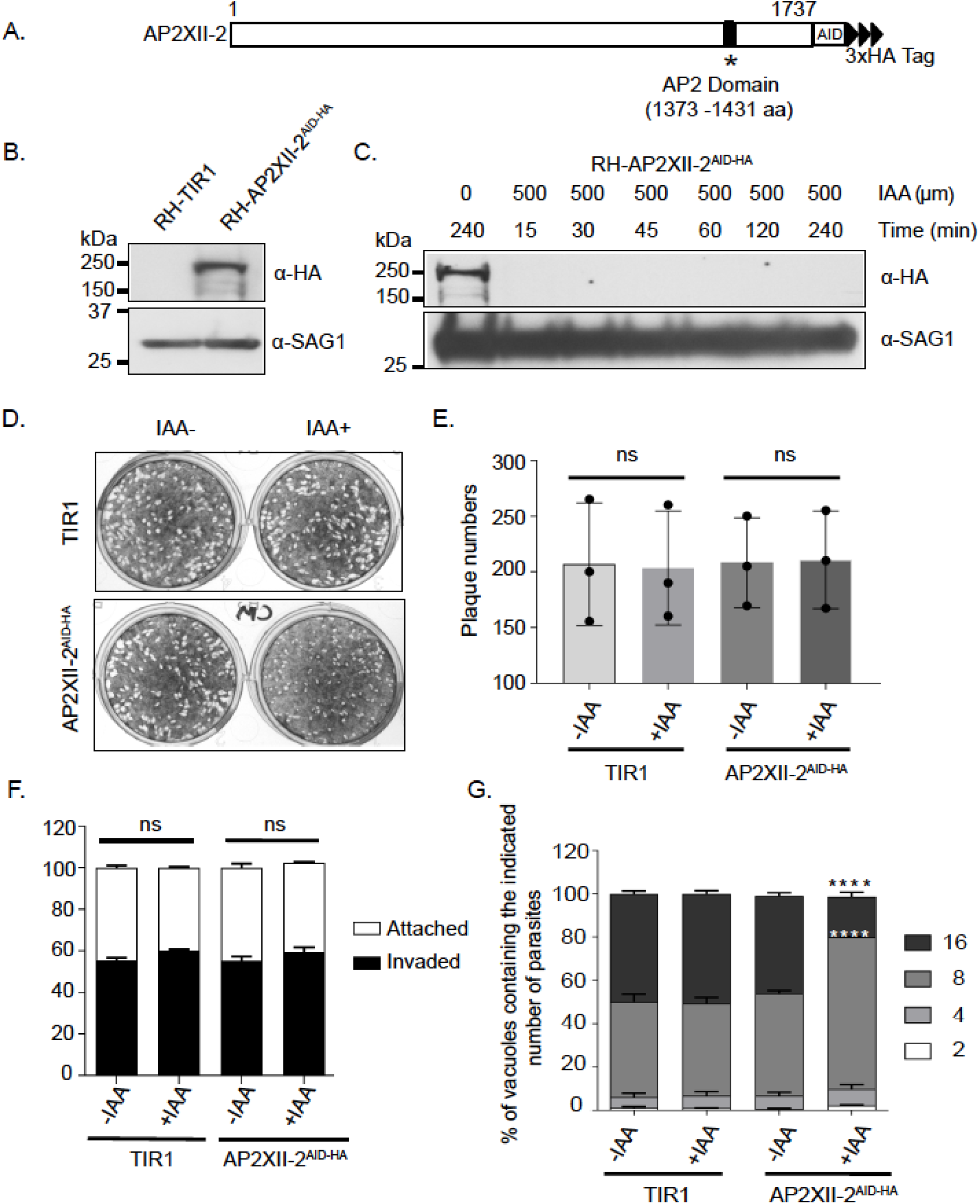
Conditional knockdown of AP2XII-2. **(A)** Schematic representation of AP2XII-2 endogenously tagged with AID-HA at the C-terminus. **(B)** Western blot showing the expression of AP2XII-2^AID-HA^ in RH-TIR1 parental parasites at the expected size (∼250kDa). SAG1 serves as a loading control. **(C)** Western blot probed with α-HA to monitor AP2XII-2^AID-HA^ protein levels over a time course after addition of 500 µM IAA. SAG1 was used as a loading control. **(D)** Plaque assay of AP2XII-2^AID-HA^ parasites treated with 500 µM IAA or vehicle (0.1% ethanol) for 6 days, showing smaller plaque sizes when AP2XII-2 is depleted. **(E)** Quantification of plaque numbers. Three independent experiments produced similar results; the data shown here represent one of the experiments (ns = not significant). **(F)** AP2XII-2^AID-HA^ parasites were grown in the presence of 500 µM IAA or vehicle for 24 hours, forcibly lysed from host cells, then allowed to invade a new HFF monolayer for 30 minutes for attachment and invasion assay (ns = not significant). **(G)** Doubling assays were performed to assess the replication of AP2XII-2_AID-HA_ parasites in the presence of 500 µM IAA or vehicle. Parasites were counted from 100 random vacuoles following 24 hours of infection. n=3 for control and 500 µM IAA treated parasites. The number of parasites were plotted as percentage of the total number of parasite vacuoles. **** denotes p<0.0001.

### Loss of AP2XII-2 delays S-phase and increases bradyzoite differentiation

We further examined the replication defect seen in AP2XII-2 depleted parasites by centrosome counting. Normally in an asynchronous parasite population in culture, 50% of parasites are in G1-phase (single centrosome) and 50% of parasites are in S-phase undergoing DNA replication (duplicated centrosome) (28). Without IAA, AP2XII-2^AID-HA^ parasites showed a 50/50 split of single and duplicated centrosomes within the population (Fig. 6). In contrast, when AP2XII-2^AID-HA^ parasites are treated with IAA, ∼75% of the population exhibited duplicated centrosomes, supporting a delay in S-phase of cell cycle. It has been shown previously that the initiation of bradyzoite differentiation and tissue cyst formation requires a slowing in DNA replication, delayed S-phase, and lengthening of G1-phase (29,30). Given the delay in S-phase produced by depletion of AP2XII-2, we examined whether there is also an increased frequency of bradyzoite development. To do so, we endogenously tagged AP2XII-2 with AID-HA in Type II ME49-TIR1 parasites (31). A clone was isolated in which AP2XII-2^AID-HA^ protein was undetectable after inclusion of IAA in the culture medium (Fig. 7A). ME49 AP2XII-2^AID-HA^ parasites treated with IAA showed the same deficiency in plaque size as seen for its RH counterpart (data not shown). We then examined the frequency of tissue cyst formation of ME49 AP2XII-2^AID-HA^ parasites incubated in alkaline stress with or without IAA (Fig. 7B). After 3 days in alkaline media without IAA, ∼50% of parasite vacuoles were staining positive with the cyst wall marker *Dolichos*. AP2XII-2^AID-HA^ parasites in IAA, however, displayed, ∼88% conversion into bradyzoite cysts (Fig. 7B-C). Type II ME49-TIR1 parasites did not show any difference in differentiation when incubated in the presence of IAA under alkaline stress conditions (data not shown) as previously “performed in (31).” It should be noted that IAA did not enhance bradyzoite formation at days 1 or 2, supporting that IAA is not a contributing factor in the increased frequency of tissue cyst formation.

**Figure 6.**
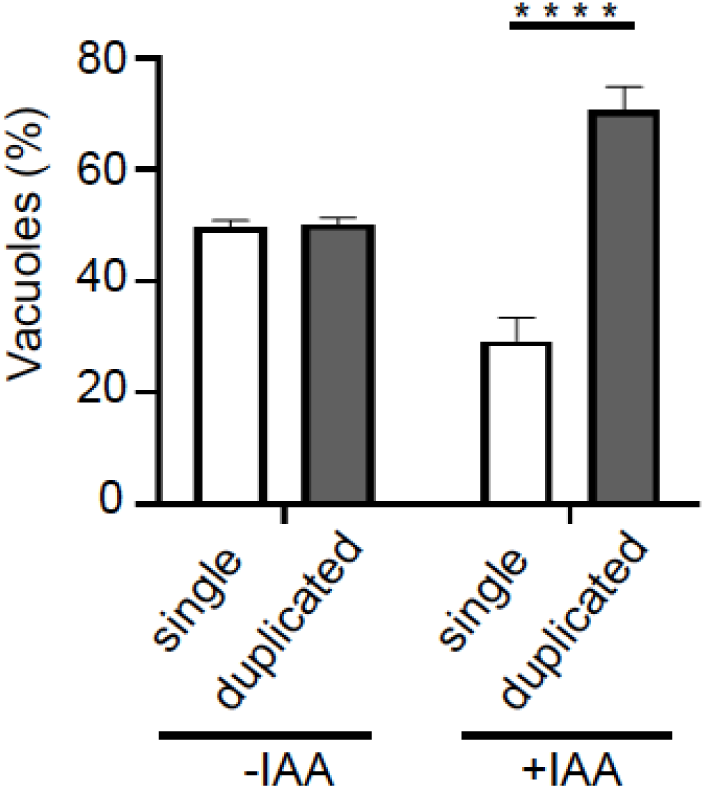
AP2XII-2 depletion results in a delay in S-phase of cell cycle. 100 parasite vacuoles in 10 random fields were counted for single or duplicated centrosomes in 3 biological replicates. Error bars represent standard deviation of the mean; **** denotes p<0.0001. Multiple comparisons were performed with two-way ANOVA.

**Figure 7.**
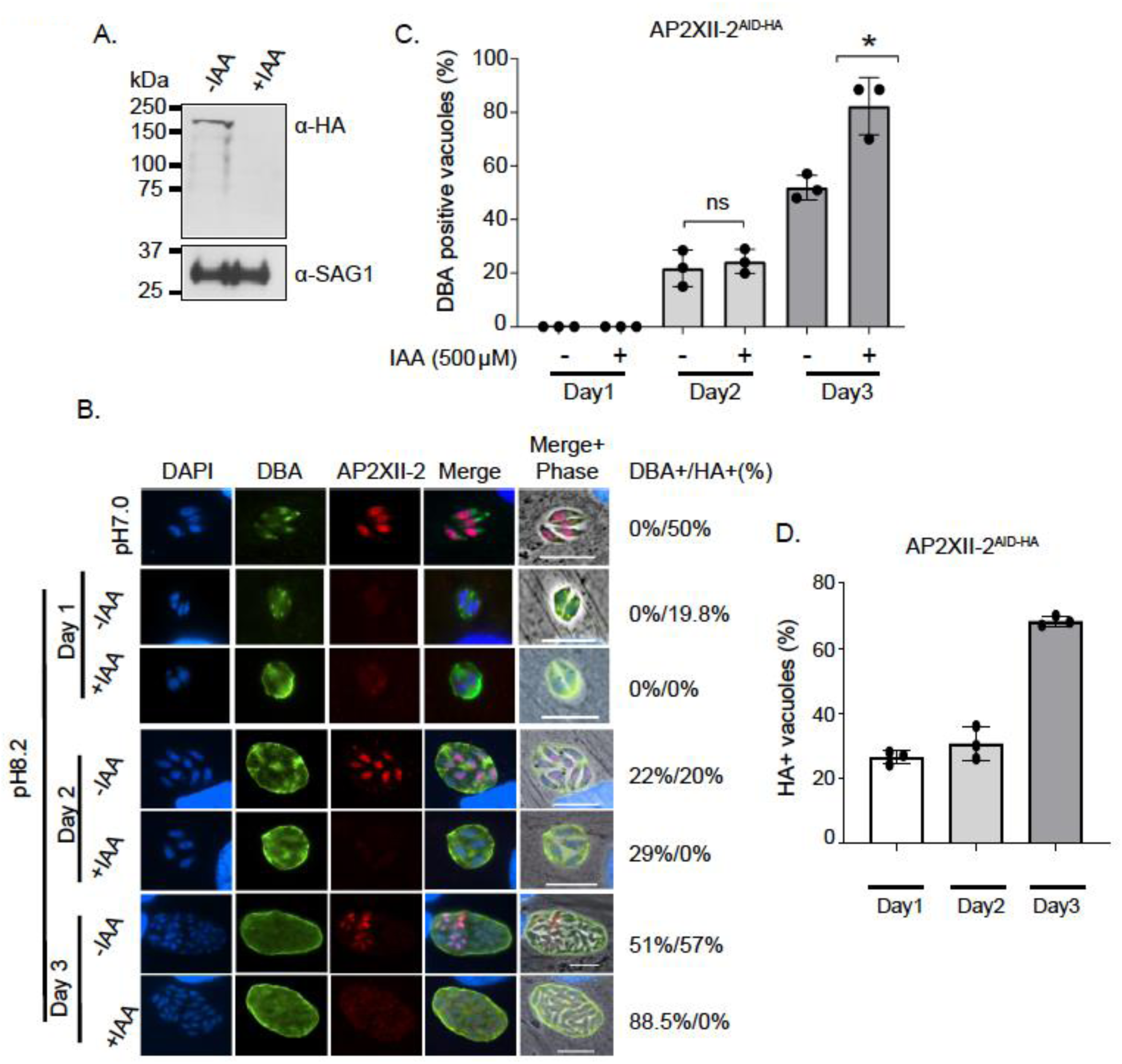
AP2XII-2 depletion leads to increased bradyzoite differentiation. **(A)** Western blot with α-HA confirms the depletion of AP2XII-2^AID-HA^ in Type II ME49-TIR1 parasites. SAG1 was used as a loading control. **(B)** Type II AP2XII-2^AID-HA^ tachyzoites were induced to differentiation with alkaline pH 8.2 for three days in the presence of 500 µM IAA or vehicle. Differentiating parasites were stained with α-HA (red) and *Dolichos* lectin (DBA, green) to visualize tissue cyst wall. DAPI (blue) was included to highlight nuclei. Scale bar = 10µM. **(C-D)** Graphical depiction of data from 7B. 100 random vacuoles were counted for *Dolichos* (DBA) and HA-positive staining, shown to the right of the IFA panels. n=3 for control and 500 µM IAA treated parasites. Error bars represent standard error of the mean; ns = not significant, * denotes p<0.05 (unpaired two-tailed student’s *t* test).

Consistent with the restricted cell cycle expression of AP2XII-2 in tachyzoites, this factor was only detected in replicating parasites following alkaline media stress (Fig. 7B), an expression pattern also seen for its interacting AP2, AP2IX-4 (23). Also similar to AP2IX-4, AP2XII-2 expression levels decrease during the initial stages of bradyzoite differentiation and then increase again. Upon addition of alkaline stress, AP2XII-2 expression levels drop from 50% to 20% within the parasite population, but by day 3, AP2XII-2 was present in ∼55% of bradyzoite cysts (Fig. 7D).

Together, these results indicate that AP2XII-2 is an important contributor for timely progression through S-phase of the cell cycle, and its loss slows tachyzoite replication and promotes bradyzoite differentiation.

## Discussion

In the present work, we report that AP2IX-4 can associate with the *T. gondii* MORC transcriptional repressor complex during its expression in S/M phases of the tachyzoite cell cycle. We also characterized a previously undescribed ApiAP2 interacting with AP2IX-4, AP2XII-2, which knockdown experiments reveal to be important for proper progression through S-phase.

There is evidence that commitment to differentiation may occur during the S/M phase of the tachyzoite cell cycle (32); it requires transient slowing of growth (33), delayed S-phase, and lengthening of the G1 phase (30). Bradyzoite cyst formation is a complex developmental pathway that requires a cascade of events in which multiple ApiAP2 transcription factors appear to be involved (11). Research is just beginning to reveal the sophistication of the ApiAP2 network, which possesses both transcriptional repressors and activators. AP2IX-9, AP2IV-4, and AP2IX-4 repress transcription of bradyzoite genes and impair tissue cyst formation (20-23), whereas AP2IV-3 appears to activate transcription (21). Given their evolutionary origin, some members of the ApiAP2 family may exhibit features found in the plant kingdom, which has classified AP2 domain proteins as active and passive transcription factors (34). Passive repressors may compete with activators to dampen transcription, allowing a fine-tuning of gene expression (34). In addition to ApiAP2s, a “master regulator” transcription factor termed BFD1 (Bradyzoite Formation Deficient-1), which harbors a myb-like DNA-binding domain, has recently been described that is essential for tissue cyst formation (35). If and how ApiAP2s interplay with BFD1 remains to be resolved.

Our study also adds to the growing evidence that at least some ApiAP2s coordinate transcriptional regulatory activities through the interaction with chromatin remodeling machinery. We recently described that the essential histone acetyltransferase (HAT) GCN5b operates in a multi-subunit complex with at least two ApiAP2 factors, AP2IX-7 and AP2XII-4, to activate gene expression (36). In the current study, we found that AP2IX-4 and APXII-2 interact with the recently described MORC complex, which contains transcriptional repressive proteins including histone deacetylase HDAC3 (24). Identifying AP2IX-4 as an interactor with MORC validates our previous observation that loss of AP2IX-4 results in upregulation of a subset of bradyzoite genes (23). It is likely that other ApiAP2s acting as repressors or activators mediate these regulatory events through collaboration with HDACs and HATs, respectively.

AP2IX-4 is a cell cycle regulated factor with peak expression in S/M phase (23). The AP2IX-4 complex we isolated contained two additional uncharacterized ApiAP2s, AP2XII-2 and AP2VIIa-3. AP2XII-2 shares a highly similar cell cycle expression pattern with AP2IX-4, which we confirmed by IFA after tagging the endogenous protein. Like AP2IX-4, AP2XII-2 is a nuclear protein with peak expression during the S/M phase. However, AP2XII-2 expression decreases before the onset of C-phase whereas AP2IX-4 remains through cytokinesis. These findings suggest that both AP2s may collaborate during tachyzoite division, but AP2IX-4 may have another independent role towards the end. Curiously, AP2IX-4 is dispensable in tachyzoites but loss of its interacting partner AP2XII-2 impairs replication due to a delay in S-phase, which likely explains the enhanced bradyzoite differentiation seen when AP2XII-2 is knocked down. AP2XII-2 may be able to compensate for the loss of AP2IX-4, but AP2IX-4 cannot replace the function of AP2XII-2. It is also possible that AP2XII-2 has an additional function in the cell cycle independent of AP2IX-4. The precise role of AP2XII-2 in facilitating progress through S-phase is an important subject for future investigation.

As we observed for AP2IX-4 (23), expression levels of AP2XII-2 protein drop during the early stages of bradyzoite differentiation. AP2XII-2 is detectable in ∼20% of bradyzoite cysts at day 1 and 2 post-induction, but is present in >55% of bradyzoite cysts by day 3 (Fig. 7B). Since the S/M period becomes more pronounced during bradyzoite differentiation, the increase in AP2XII-2 positive parasites further supports a defined cell cycle context for commitment to bradyzoite differentiation. Previously, we found that AP2IX-4 is present only in dividing parasites within cysts, and this is likely to be the case for AP2XII-2. We suspect these in vitro bradyzoites are immature and still undergoing some level of cell division, which requires these two ApiAP2s for proper progression through the cell cycle. Our findings underscore the link between cell cycle progression, parasite division, and the commitment to bradyzoite differentiation, and bolster the concept that cell cycle regulated ApiAP2s like AP2IX-4 and AP2XII-2 work with with chromatin remodeling machinery like MORC to regulate gene expression relevant to cell cycle progression.

## Materials and Methods

### Parasite culture and transfection

Parasite strains used for this study include RHΔ*hxgprt*Δ*ku80* (37), RHΔ*hxgprt*Δ*ku80*:*AP2IX-4*^*HA*^ (23), and RH and ME49 parasites engineered to express TIR1 for IAA-mediated protein degradation (27, 31).

Parasites were cultured in human foreskin fibroblast cells (HFF) in Dulbecco’s modified eagle medium supplemented with 1% fetal bovine serum, 100 unit/ml penicillin and 100 µg/ml streptomycin. Transfection of parasites was performed as previously described using the 4D-Nucleofector System (Lonza) and selected for drug-resistance 24 hrs post-transfection (38-40). Transfected parasites were selected in either 1 µM pyrimethamine or 25 µg/ml mycophenolic acid + 50 µg/ml xanthine as described (41,42).

### Generation of parasites expressing endogenously tagged AP2IX-4^HA^ and AP2XII-2^MYC^

RHΔ*hxgprt*Δ*ku80*:*AP2IX-4*^*HA*^ (23) parasites were genetically modified to express endogenous AP2XII-2 tagged with MYC at the C-terminus. Modification of the endogenous locus of AP2XII-2 was performed by a genetic knock-in approach using plasmid pLIC-3xMYC-HXGPRT (37). Primers (Fw 5’ ATCCAATTTAATTAATCTTCTGTGTCCCGGTGG and Rv 5’ TCCAATTTTAATTAAGCCACTGAGTGGTGAAACA 3’) were used to amplify 1,975 bp of the AP2XII-2 gene, which was then cloned into the PacI site of pLIC-3xMYC-HXGPRT plasmid. The resulting construct was linearized and transfected into RHΔ*hxgprt*Δ*ku80:AP2IX-4*^*HA*^ parasites. Transfected parasites were selected in 25 µg/ml mycophenolic acid + 50 µg/ml xanthine and independent clones were isolated by limiting dilution.

### Generation of parasites expressing endogenously tagged AP2XII-2^AID-HA^

To generate Type I and Type II conditional knock-down strains, specific primers (Fw 5’ ACCGGGCCCGCTAGCTCTTCTGTGTCCCGGTGG 3’ and Rv 5’ CGAGCCCTTGCTAGCGCCACTGAGTGGTGAAACA 3’) were used to amplify 1,975 bp of the AP2XII-2 gene for cloning into the Nhe1 site of pLIC-AID-3xHA-DHFR plasmid. The construct was linearized and transfected into either RH or ME49 TIR1-expressing parasites (27,31). Parasites were selected in 1 µM pyrimethamine and single clones were isolated by limiting dilution. Parasites were treated with 500 µM IAA (Sigma-Aldrich) or vehicle for indicated period of time to induce degradation of AID-tagged protein.

### Parasite growth assays

For plaque assays, 500 freshly syringe-lysed intracellular parasites were allowed to invade a confluent monolayer of HFFs grown in 12-well plates in 500 µM IAA or vehicle. Six days post-infection, the infected monolayers were stained with crystal violet stain to determine host cell lysis as previously described (43).

For doubling assays, purified intracellular parasites were allowed to invade confluent HFFs for 30 min. After invasion, infected HFFs were gently washed with culture medium to remove extracellular tachyzoites and fresh media was added. After 24 hrs, infected monolayers were fixed with 4% paraformaldehyde for 10 min at room temperature, incubated in blocking buffer for 30 min, then stained with α-SAG1 antibody (Invitrogen) for 1 hr at room temperature. The fixed cells were then washed with PBS and incubated with secondary goat anti-mouse Alexa-fluor 488 (Invitrogen) for 1 hr at room temperature. Cells were mounted with ProLong Gold Antifade Mounting solution (Invitrogen) containing DAPI. The number of parasites in 100 randomly selected vacuoles was counted at the designated time point (44).

### Invasion and attachment assay

Invasion and attachment assays were performed as previously described (45). Briefly, parasites were allowed to grow in 500 µM IAA or vehicle for 24 hrs, then purified from host cells via syringe lysis. The parasites were allowed to invade confluent HFFs for 30 min prior to fixation with 4% paraformaldehyde (Sigma). Attached parasites were labeled with mouse α-SAG1 (Thermo Fisher) and then subjected to permeabilization in 0.1% Triton X-100. Next, parasites were stained with α-IMC3 antibody (a gift from Dr. Marc-Jan Gubbels) followed by α-mouse Alexa-fluor 488 (Invitrogen) and α-rat Alexa-fluor 594 (Invitrogen). Ten random fields were imaged and the number of attached and invaded parasites was counted. Primary antibody dilutions were used as follows: mouse α-SAG1 (Thermo Fisher) 1:1,000 and rat α-IMC3 1:2,000. Secondary Alexa fluor antibodies were used at 1:1,000 dilution.

### Western blotting

Confluent HFFs infected with tachyzoites were syringe-lysed 24 hrs post-infection and filtered, followed by centrifugation. Parasite pellets were resuspended in NuPAGE lysis buffer and boiled for 5 min. Eighty µg of parasite lysates were separated on a 4–12% Tris-acetate polyacrylamide gradient gel (Invitrogen) and transferred onto PVDF membranes. Western blotting was performed with the designated antibodies and developed with SuperSignal West Femto Sensitivity Substrate (Pierce).

Primary antibody dilutions were used as follows: rat α-HA antibody (Roche) 1:2,000; mouse α-MYC (Cell Signaling) 1:5,000; rabbit α-MYC (Thermo) 1:5,000; rabbit α-H3 (Sigma) 1:5,000; rabbit eIF2α 1:20,000 (46), mouse α-Sag1 (Thermo Fisher) 1:5,000. Secondary horseradish peroxidase (HRP)-conjugated antibody dilutions were used as follows: goat α-rat (GE Healthcare) 1:2,000; goat α-mouse (Sigma) 1:5,000; goat α-rabbit (Sigma) 1:5,000.

### Immunofluorescence assays (IFA)

Parasites growing in confluent HFFs were fixed in 4% paraformaldehyde for 10 min followed by washing in PBS three times. Fixed cells were incubated for 1 hr at room temperature in blocking buffer containing 3% BSA and 0.2% Triton X-100. The cells were incubated with the designated antibodies in blocking buffer at 4°C overnight followed by washing in PBS. The cells were then incubated with secondary antibodies coupled to Alexa 488/594/568/647 at room temperature for 1 hr. The cells were finally washed with PBS and mounted with ProLong Gold Antifade Mounting solution (Invitrogen) containing DAPI (Invitrogen) and then visualized using a Nikon Eclipse E100080i microscope. Images were captured with a Hamamatsu C4742-95 CCD camera. Nikon NIS element software was used to analyze and capture images. Primary antibody dilutions were used as followed: rabbit α-HA (Cell Signaling) 1:1,000, rat α-HA (Roche), mouse α-MYC (Cell Signaling) 1:1,000, *Toxoplasma* α-Centrin-1 (Kerafast company) 1:2,000, rat α-IMC3 (supplied by Dr. Marc-Jan Gubbels) 1:2,000. For the visualization of brazyzoite tissue cyst walls, FITC-conjugated *Dolichos biflorus* lectin (Vector Laboratories) was used at a 1:500 dilution for 1 hr at room temperature as previously described (23).

### Bradyzoite differentiation assay

*In vitro* bradyzoite differentiation experiments were performed as described elsewhere (20,47,48). Briefly, parasites were allowed to invade a confluent monolayer of HFFs grown in 12-well plates. After 2 hrs post-invasion, tachyzoite media (pH 7.0) was replaced with bradyzoite differentiation media (pH 8.2), and parasites were grown in the presence of IAA (500 µM) or vehicle for indicated time periods (20,47,48). Media was changed every 24 hrs to maintain pH for parasites undergoing differentiation, with fresh IAA or vehicle added.

### Subcellular fractionation

Purified parasite pellets were resuspended in 1 ml low salt buffer (50 mM HEPES-NaOH pH 7.5, 20% glycerol, 10 mM NaCl, 0.1% NP40) and incubated at 4°C for 10 min followed by centrifugation at 2,500x*g* for 15 min. The supernatant was collected as the cytoplasmic fraction. The nuclei-containing pellet was resuspended in high salt extraction buffer (50 mM HEPES-NaOH pH 7.5, 20% glycerol, 420 mM NaCl, 0.4% NP40) for 10 min at 4°C with intermittent stirring followed by brief sonication (5 rounds, 10 seconds each). Following centrifugation at 13,000x*g* for 30 min, the cleared nuclear fraction supernatant was collected. All buffers were supplemented with (Roche – Complete Mini, EDTA-free, Cat. no. 11 836 170 001).

### Purification of the AP2IX-4 complex

RHΔ*hxgprt* parasites endogenously expressing AP2IX-4^HA^ or AP2IX-4^HA^/AP2XII-2^MYC^ were used to pull-down the core AP2IX-4 complex. Large-scale tachyzoite cultures were grown for 24 hrs in HFFs, followed by scraping the infected monolayers and syringe-lysing through a 25-gauge needle to isolate free tachyzoites. Parasites were further purified through a 3 micron membrane filter and centrifuged at 4°C for at 10 min at 700x*g*. The parasite pellets were lysed in low salt buffer (10 mM HEPES pH 7.4, 10 mM KCl, 10% [v/v] glycerol, 0.1% [v/v] NP-40) for 10 min on ice followed by centrifugation at 2,500x*g* for 10 min to isolate soluble cytosolic and nuclear pellet fractions. The nuclear pellet was washed twice with low salt buffer and then treated with MNase (Thermo Fisher Scientific) in the presence of 1 mM CaCl_2_ for 10 min at 37°C followed by adding EDTA to stop MNase digestion. The samples were centrifuged again to isolate MNAse soluble chromatin fraction and MNAse insoluble pellet fractions. The MNAse insoluble pellet fractions was dissolved in high salt buffer (10 mM HEPES pH 7.4, 10 mM KCl, 10% [v/v] glycerol, 0.1% [v/v] NP-40) containing 420 mM NaCl for 30 min while rocking at 4°C. Finally, the high salt fraction was isolated by centrifugation. Soluble fractions were combined for immunoprecipitation with α-HA or α-MYC antibodies coupled with Magnetic Dynabeads as per manufacturer’s instructions (Thermo Fisher). Immunoprecipitated proteins on the beads were boiled in SDS sample buffer for 5 min and processed for western blotting or proteomics analysis. All buffers used in the protocol were supplemented with protease inhibitor cocktail mixture (Roche – Complete Mini, EDTA-free, Cat. no. 11 836 170 001) and 1mM PMSF (Roche).

### Mass spectrometry of the AP2IX-4 complex

Sample preparation and mass spectrometry analysis were performed as previously described (36). Briefly, immunoprecipitated samples on Dynabeads were resuspended in 8M Urea for denaturation, reduced, and alkylated. Alkylated samples were digested with endoproteinase LysC and Trypsin gold (Promega) at 37°C overnight. Digested peptides were purified with a spin column (Pierce), followed by injection onto a C18 3 µM reversed phase trap column. The peptides were then separated on an EASY-Spray 2 µM 15 cm reversed phase analytical column (Thermo Fisher). Peptides were separated on acetonitrile (2-25% gradient for 90 min) in front of a Velos Pro Orbitrap mass spectrometer in data-dependent acquisition mode. Bioinformatics analysis was performed using a custom *T. gondii* database as described in (17). SAINT (Significance Analysis of INTeractome) analysis was performed to analyze interactome data as previously described (36).

### Statistical analysis

GraphPad Prism software was used for statistical analyses. SAINTexpress software was used for SAINT analysis.

## Acknowledgements

This research was supported by National Institutes of Health grants AI124682 (M.W.W.) and AI116496 (W.J.S.). The authors thank Dr. Marc-Jan Gubbels (Boston College) for supplying rat α-IMC3 antibody and Dr. David Sibley (Washington University) for supplying the TIR1-expressing parasite lines.

## Figure Legends

**Supplemental Table 1**. Complete proteomics dataset for mass spectrometry analysis of AP2XII-2 immunoprecipitate.

## References

1. Dubey JP, Miller NL, Frenkel JK. 1970. Toxoplasma gondii life cycle in cats. J Am Vet Med Assoc 157:1767–70.

2. Dubey JP, Hill DE, Jones JL, Hightower AW, Kirkland E, Roberts JM, Marcet PL, Lehmann T, Vianna MC, Miska K, Sreekumar C, Kwok OC, Shen SK, Gamble HR. 2005. Prevalence of viable Toxoplasma gondii in beef, chicken, and pork from retail meat stores in the United States: risk assessment to consumers. J Parasitol 91:1082–93.

3. Jones JL, Kruszon-Moran D, Wilson M, McQuillan G, Navin T, McAuley JB. 2001. Toxoplasma gondii infection in the United States: seroprevalence and risk factors. Am J Epidemiol 154:357–65.

4. Martinez AJ, Sell M, Mitrovics T, Stoltenburg-Didinger G, Iglesias-Rozas JR, Giraldo-Velasquez MA, Gosztonyi G, Schneider V, Cervos-Navarro J. 1995. The neuropathology and epidemiology of AIDS. A Berlin experience. A review of 200 cases. Pathol Res Pract 191:427–43.

5. Tenter AM, Heckeroth AR, Weiss LM. 2000. Toxoplasma gondii: from animals to humans. Int J Parasitol 30:1217–58.

6. Jones J, Lopez A, Wilson M. 2003. Congenital toxoplasmosis. Am Fam Physician 67:2131–8.

7. Ortiz-Alegria LB, Caballero-Ortega H, Canedo-Solares I, Rico-Torres CP, Sahagun-Ruiz A, Medina-Escutia ME, Correa D. 2010. Congenital toxoplasmosis: candidate host immune genes relevant for vertical transmission and pathogenesis. Genes Immun 11:363–73.

8. Murata Y, Sugi T, Weiss LM, Kato K. 2017. Identification of compounds that suppress Toxoplasma gondii tachyzoites and bradyzoites. PLoS One 12:e0178203.

9. Montazeri M, Mehrzadi S, Sharif M, Sarvi S, Shahdin S, Daryani A. 2018. Activities of anti-Toxoplasma drugs and compounds against tissue cysts in the last three decades (1987 to 2017), a systematic review. Parasitol Res 117:3045–3057.

10. Benmerzouga I, Checkley LA, Ferdig MT, Arrizabalaga G, Wek RC, Sullivan WJ, Jr. 2015. Guanabenz repurposed as an antiparasitic with activity against acute and latent toxoplasmosis. Antimicrob Agents Chemother 59:6939–45.

11. Jeffers V, Tampaki Z, Kim K, Sullivan WJ, Jr. 2018. A latent ability to persist: differentiation in Toxoplasma gondii. Cell Mol Life Sci 75:2355–2373.

12. Skariah S, McIntyre MK, Mordue DG. 2010. Toxoplasma gondii: determinants of tachyzoite to bradyzoite conversion. Parasitol Res 107:253–60.

13. Sullivan WJ, Jr. 2020. Mastering Toxoplasma sex and sleep. Nat Microbiol 5:533–534.

14. Balaji S, Babu MM, Iyer LM, Aravind L. 2005. Discovery of the principal specific transcription factors of Apicomplexa and their implication for the evolution of the AP2-integrase DNA binding domains. Nucleic Acids Res 33:3994–4006.

15. Altschul SF, Wootton JC, Zaslavsky E, Yu YK. 2010. The construction and use of log-odds substitution scores for multiple sequence alignment. PLoS Comput Biol 6:e1000852.

16. Behnke MS, Wootton JC, Lehmann MM, Radke JB, Lucas O, Nawas J, Sibley LD, White MW. 2010. Coordinated progression through two subtranscriptomes underlies the tachyzoite cycle of Toxoplasma gondii. PLoS One 5:e12354.

17. Gajria B, Bahl A, Brestelli J, Dommer J, Fischer S, Gao X, Heiges M, Iodice J, Kissinger JC, Mackey AJ, Pinney DF, Roos DS, Stoeckert CJ, Jr., Wang H, Brunk BP. 2008. ToxoDB: an integrated Toxoplasma gondii database resource. Nucleic Acids Res 36:D553–6.

18. Kafsack BF, Rovira-Graells N, Clark TG, Bancells C, Crowley VM, Campino SG, Williams AE, Drought LG, Kwiatkowski DP, Baker DA, Cortes A, Llinas M. 2014. A transcriptional switch underlies commitment to sexual development in malaria parasites. Nature 507:248–52.

19. Yuda M, Iwanaga S, Shigenobu S, Kato T, Kaneko I. 2010. Transcription factor AP2-Sp and its target genes in malarial sporozoites. Mol Microbiol 75:854–63.

20. Radke JB, Lucas O, De Silva EK, Ma Y, Sullivan WJ, Jr., Weiss LM, Llinas M, White MW. 2013. ApiAP2 transcription factor restricts development of the Toxoplasma tissue cyst. Proc Natl Acad Sci U S A 110:6871–6.

21. Hong DP, Radke JB, White MW. 2017. Opposing Transcriptional Mechanisms Regulate Toxoplasma Development. mSphere 2.

22. Radke JB, Worth D, Hong D, Huang S, Sullivan WJ, Jr., Wilson EH, White MW. 2018. Transcriptional repression by ApiAP2 factors is central to chronic toxoplasmosis. PLoS Pathog 14:e1007035.

23. Huang S, Holmes MJ, Radke JB, Hong DP, Liu TK, White MW, Sullivan WJ, Jr. 2017. Toxoplasma gondii AP2IX-4 Regulates Gene Expression during Bradyzoite Development. mSphere 2.

24. Farhat DC, Swale C, Dard C, Cannella D, Ortet P, Barakat M, Sindikubwabo F, Belmudes L, De Bock PJ, Coute Y, Bougdour A, Hakimi MA. 2020. A MORC-driven transcriptional switch controls Toxoplasma developmental trajectories and sexual commitment. Nat Microbiol 5:570–583.

25. Gubbels MJ, Wieffer M, Striepen B. 2004. Fluorescent protein tagging in Toxoplasma gondii: identification of a novel inner membrane complex component conserved among Apicomplexa. Mol Biochem Parasitol 137:99–110.

26. Fung C, Beck JR, Robertson SD, Gubbels MJ, Bradley PJ. 2012. Toxoplasma ISP4 is a central IMC sub-compartment protein whose localization depends on palmitoylation but not myristoylation. Mol Biochem Parasitol 184:99–108.

27. Brown KM, Long S, Sibley LD. 2017. Plasma Membrane Association by N-Acylation Governs PKG Function in Toxoplasma gondii. mBio 8.

28. Alvarez CA, Suvorova ES. 2017. Checkpoints of apicomplexan cell division identified in Toxoplasma gondii. PLoS Pathog 13:e1006483.

29. Jerome ME, Radke JR, Bohne W, Roos DS, White MW. 1998. Toxoplasma gondii bradyzoites form spontaneously during sporozoite-initiated development. Infect Immun 66:4838–44.

30. Radke JR, Guerini MN, Jerome M, White MW. 2003. A change in the premitotic period of the cell cycle is associated with bradyzoite differentiation in Toxoplasma gondii. Mol Biochem Parasitol 131:119–27.

31. Brown KM, Sibley LD. 2018. Essential cGMP Signaling in Toxoplasma Is Initiated by a Hybrid P-Type ATPase-Guanylate Cyclase. Cell Host Microbe 24:804–816 e6.

32. White MW, Radke JR, Radke JB. 2014. Toxoplasma development - turn the switch on or off? Cell Microbiol 16:466–72.

33. Bohne W, Heesemann J, Gross U. 1994. Reduced replication of Toxoplasma gondii is necessary for induction of bradyzoite-specific antigens: a possible role for nitric oxide in triggering stage conversion. Infect Immun 62:1761–7.

34. Licausi F, Ohme-Takagi M, Perata P. 2013. APETALA2/Ethylene Responsive Factor (AP2/ERF) transcription factors: mediators of stress responses and developmental programs. New Phytol 199:639–49.

35. Waldman BS, Schwarz D, Wadsworth MH, 2nd, Saeij JP, Shalek AK, Lourido S. 2020. Identification of a Master Regulator of Differentiation in Toxoplasma. Cell 180:359–372 e16.

36. Harris MT, Jeffers V, Martynowicz J, True JD, Mosley AL, Sullivan WJ, Jr. 2019. A novel GCN5b lysine acetyltransferase complex associates with distinct transcription factors in the protozoan parasite Toxoplasma gondii. Mol Biochem Parasitol 232:111203.

37. Huynh MH and Carruthers VB. 2009. Tagging of endogenous genes in a Toxoplasma gondii strain lacking Ku80. Eukaryot Cell 8(4): 530–539.

38. Harding CR, Egarter S, Gow M, Jimenez-Ruiz E, Ferguson DJ, Meissner M. 2016. Gliding Associated Proteins Play Essential Roles during the Formation of the Inner Membrane Complex of Toxoplasma gondii. PLoS Pathog 12:e1005403.

39. Upadhya R, Kim K, Hogue-Angeletti R, Weiss LM. 2011. Improved techniques for endogenous epitope tagging and gene deletion in Toxoplasma gondii. J Microbiol Methods 85:103–13.

40. Soldati D, Boothroyd JC. 1993. Transient transfection and expression in the obligate intracellular parasite Toxoplasma gondii. Science 260:349–52.

41. Donald RG, Roos DS. 1993. Stable molecular transformation of Toxoplasma gondii: a selectable dihydrofolate reductase-thymidylate synthase marker based on drug-resistance mutations in malaria. Proc Natl Acad Sci U S A 90:11703–7.

42. Donald RG, Carter D, Ullman B, Roos DS. 1996. Insertional tagging, cloning, and expression of the Toxoplasma gondii hypoxanthine-xanthine-guanine phosphoribosyltransferase gene. Use as a selectable marker for stable transformation. J Biol Chem 271:14010–9.

43. Roos DS, Donald RG, Morrissette NS, Moulton AL. 1994. Molecular tools for genetic dissection of the protozoan parasite Toxoplasma gondii. Methods Cell Biol 45:27–63.

44. Fichera ME, Bhopale MK, Roos DS. 1995. In vitro assays elucidate peculiar kinetics of clindamycin action against Toxoplasma gondii. Antimicrob Agents Chemother 39:1530–7.

45. Huynh MH, Rabenau KE, Harper JM, Beatty WL, Sibley LD, Carruthers VB. 2003. Rapid invasion of host cells by Toxoplasma requires secretion of the MIC2-M2AP adhesive protein complex. EMBO J 22:2082–90.

46. Augusto L, Martynowicz J, Staschke KA, Wek RC, Sullivan WJ, Jr. 2018. Effects of PERK eIF2alpha Kinase Inhibitor against Toxoplasma gondii. Antimicrob Agents Chemother 62:11.

47. Bohne W, Hunter CA, White MW, Ferguson DJ, Gross U, Roos DS. 1998. Targeted disruption of the bradyzoite-specific gene BAG1 does not prevent tissue cyst formation in Toxoplasma gondii. Mol Biochem Parasitol 92:291–301.

48. Weiss LM, Laplace D, Takvorian PM, Tanowitz HB, Cali A, Wittner M. 1995. A cell culture system for study of the development of Toxoplasma gondii bradyzoites. J Eukaryot Microbiol 42:150–7.

